# RFMix-reader: Accelerated reading and processing for local ancestry studies

**DOI:** 10.1101/2024.07.13.603370

**Authors:** Kynon J.M. Benjamin

## Abstract

**Motivation:** Local ancestry inference is a powerful technique in genetics, revealing population history and the genetic basis of diseases. It is particularly valuable for improving eQTL discovery and fine-mapping in admixed populations. Despite the widespread use of the `RFMix` software for local ancestry inference, large-scale genomic studies face challenges of high memory consumption and processing times when handling `RFMix` output files.

**Results:** Here, I present `RFMix-reader`, a new Python-based parsing software, designed to streamline the analysis of large-scale local ancestry datasets. This software prioritizes computational eiciency and memory optimization, leveraging GPUs when available for additional speed boosts. By overcoming these data processing hurdles, `RFMix-reader` empowers researchers to unlock the full potential of local ancestry data for understanding human health and health disparities.

**Availability:** `RFMix-reader` is freely available on PyPI at https://pypi.org/project/rfmix-reader/, implemented in Python 3, and supported on Linux, Windows, and Mac OS.

**Supplementary information:** Supplementary data are available at https://rfmix-reader.readthedocs.io/en/latest/.

## Introduction

Local ancestry inference, pinpointing ancestry at specific genomic regions within individuals, is a powerful technique in genetics. This method not only provides valuable insights into population history (Peter, 2016) but also holds immense potential for unraveling the genetic basis of complex traits (Zhong *et al*., 2019). Recent studies have demonstrated that local ancestry information is particularly valuable for improving eQTL (expression quantitative trait loci) discovery (Ehsan *et al*., 2024) and fine-mapping in admixed populations (Gay *et al*., 2020). While `RFMix` (Maples *et al*., 2013) is one of the most widely used software for local ancestry inference, a significant bottleneck exists for large-scale studies: the manipulation and processing of output files generated by `RFMix`. Standard scripting solutions, even when leveraging GPUs, often encounter limitations due to high memory consumption and excessive processing times.

To address these limitations, I introduce `RFMix-reader`, a Python-based parsing software specifically designed to streamline the analysis of local ancestry data. `RFMix-reader` prioritizes both computational eiciency and memory optimization, enabling researchers to work effectively with large datasets common in eQTL analysis studies. Its capability to leverage GPUs, in addition to eicient CPU processing, offers substantial speed boosts. By overcoming these data processing hurdles, `RFMix-reader` empowers researchers to more effectively use local ancestry information, leading to a deeper understanding of the genetic underpinnings of health disparities.

## Implementation

### RFMix-reader

The core functionality of `RFMix-reader` lies within the `read_rfmix` function (Figure 1 A), designed for eicient loading and processing of RFMix output files. This function takes a `file_prefix` argument to identify the relevant RFMix files and uses the `get_prefixes` function to gather the `*.fb.tsv` (local ancestry) and `*.rfmix.Q` (global ancestry) RFMix output file paths. `RFmix-reader` prioritizes eicient data handling by checking for GPU availability. If a GPU is present, it leverages `cudf` for simple DataFrame operations; otherwise, it defaults to `pandas` (The pandas development team, 2024) for broader compatibility. The function provides visual feedback during execution using `tqdm` [doi:10.5281/zenodo.595120] progress bars (enabled by default with `verbose = True`). Upon successful processing, `read_rfmix` returns three key components: loci information for local ancestry (DataFrame), global ancestry per chromosome (DataFrame), and local ancestry data (Dask Array).

**Figure 1:**
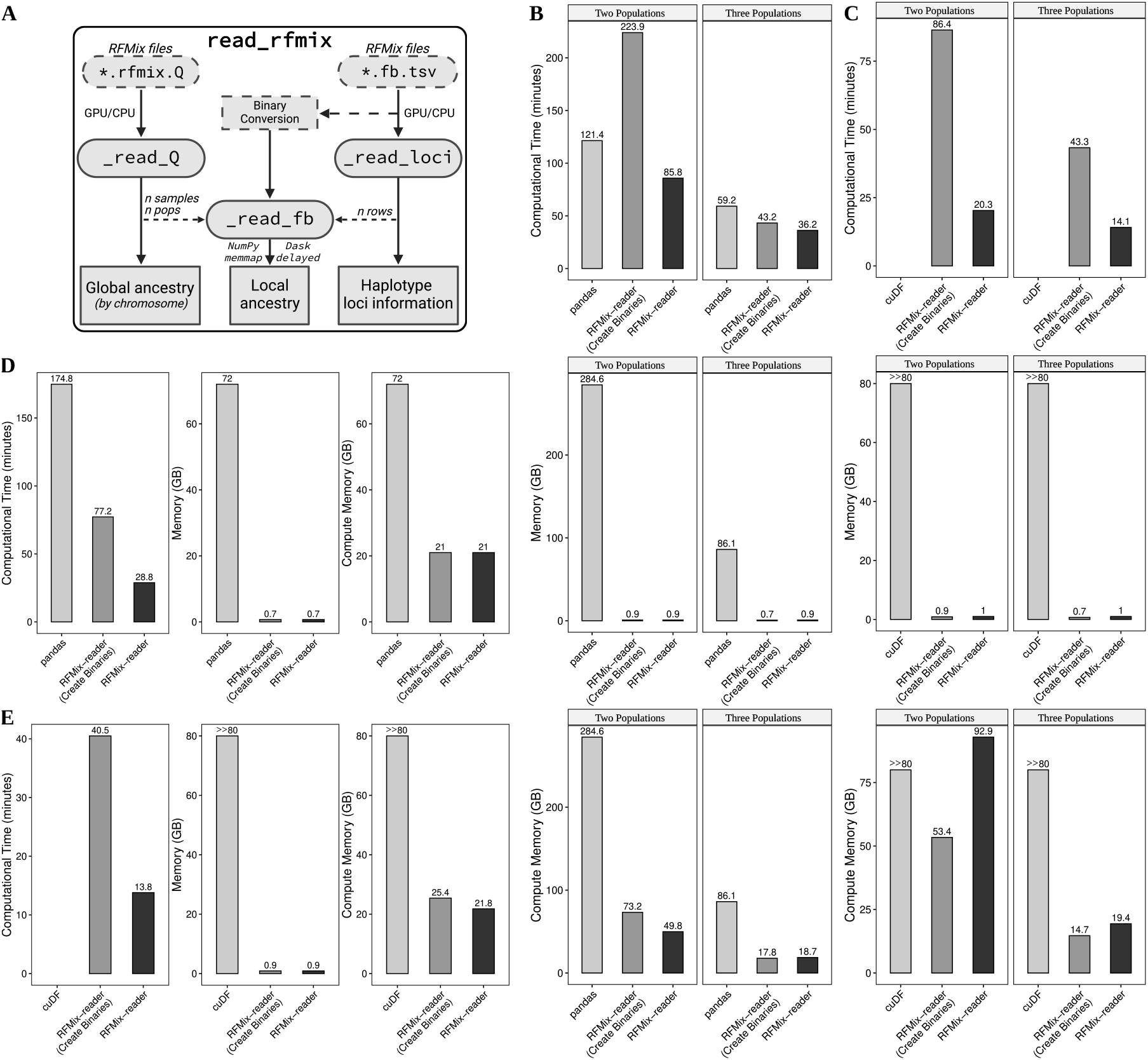
`RFMix-reader` reduces computational time and memory usage compared to `pandas` and `cuDF`. A. Flowchart of `RFMix-reader` created with BioRender.com. Bar plots of simulated two-population (n=500; 12.3M loci) and three-population (n=100; 12.3M loci) data comparing computation time (top), memory usage (middle), and memory after data computation (bottom) for B. CPU only and C. GPU available. Bar plots of real data for Black American two-population ancestry model from the African Ancestry Neuroscience Research Initiative (Benjamin *et al*., 2024) (n=526; 2.9M loci) comparing computation time (left), memory usage (middle), and memory after data computation (right) for D. CPU only and E. GPU available. Data loading failed for `cuDF` due to GPU memory limit (A100 with 80 GB of memory); as such, memory consumption (`>>80` GB) and computational time are unknown. Comparisons for `RFMix-reader` are shown with and without the generation of binary files.

### Extracting and processing RFMix data

`RFMix-reader` employs several internal helper functions for specific data-handling tasks:

- `_read_loci` : This function extracts local ancestry loci information from the `*.fb.tsv` files. For eiciency, only the “chromosome” and “physical_position” columns are loaded into memory. Additionally, the function adds a sequential index column for easier reference similar to `bim` output with `pandas-plink`.
- `_read_Q` : This function retrieves global ancestry data from `*.rfmix.Q` files. It reads the Q matrix and incorporates chromosome information extracted from the filename.
- `_read_fb` : This function tackles local ancestry data from `*.fb.tsv` files. It loads the data into a Dask Array for eicient processing of large datasets. To minimize memory usage, the function uses chunked reading with `numpy.memmap` (Harris *et al*., 2020) and Dask’s `delayed` computations, allowing it to effectively handle files exceeding available memory.

### Optimizing local ancestry data handling

`RFMix-reader` offers flexibility for handling local ancestry data in `*.fb.tsv` files. To leverage `memmap`, binary files are required. To optimize reading speed, consider generating binary versions of haplotype information from the `*.fb.tsv` files either beforehand using the separate `create_binaries` function (recommended for repeated analyses) or on the fly during `read_rfmix` by setting `generate_binary=True`. Be aware that generating binary files for a complete chromosome set can be time-consuming, potentially taking 20 minutes (or more). Additionally, `read_rfmix` uses the `_subset_populations` function internally to process the local ancestry haplotype data. This function plays a crucial role by subsetting the data and summing adjacent columns for each specific population, effectively combining haplotypes for each reference population. These considerations ensure eicient handling of large-scale local ancestry datasets.

### Benchmarking and application

To assess performance, `RFMix-reader` was compared to standard optimized scripting using `pandas` and `cuDF`. I employed two admixed simulated genotypes (two and three population admixes) generated using Haptools (Massarat *et al*., 2023) and reference populations from the 1000 Genomes Project (*et al*., 2015). For the two-population admixture, I simulated genotypes for 500 individuals across ten generations (20% Utah residents with Northern and Western European ancestry [CEU]; 80% Yoruba [YRI]; 12,317,365 loci). The three-population scenario involved 100 individuals simulated over ten generations (34% CEU; 65% YRI; 1% Puerto Rican; 12,316,809 loci). In both scenarios, `cuDF` encountered limitations due to GPU memory constraints (80 GB for A100) and was unable to process the full chromosome set. Consequently, only `RFMix-reader` could leverage the GPU for faster file reading observed through a reduction of 25% compared with CPU usage of `RFMix-reader` (Figure 1 B, C). Furthermore, `RFMix-reader` demonstrated substantial reductions in memory consumption (both with and without in-memory processing and binary file generation) and computational time compared to optimized scripting (Figure 1 B, C).

I further evaluated `RFMix-reader` using real data from the African Ancestry Neuroscience Research Initiative (Benjamin *et al*., 2024) (n=526; 2,890,613 loci). Similar to the simulated data, `cuDF` processing was hampered by GPU memory limitations (80 GB for A100) and could not handle the complete chromosome set. Here too, `RFMix-reader` exhibited substantial reductions in processing time and memory usage, particularly when a GPU was available, compared to standard optimized scripting (Figure 1 D, E). Overall, these results demonstrate the initial utility and effectiveness of `RFMix-reader` for parsing real-world datasets.

## Acknowledgments

I would like to extend my deepest appreciation to Aaqib Ansari and Karen Ives for their comments and suggestions.

## Data availability

Analysis-ready genotype data will be shared with researchers who obtain database of Genotypes and Phenotype (dbGaP) access (phs000979.v3.p2). The 1000 Genomes Project reference data are available at http://www.internationalgenome.org/data/. The simulated data used in the benchmarking analyses are available on Synapse syn61691659. The real data will be shared with researchers who obtain access to the genotype data (phs000979.v3.p2) upon request. All scripts generated in this paper are available through GitHub at https://github.com/heart-gen/rfmix_reader-benchmarking.

